# The intrinsic spatiotemporal organization of the human brain - A multi-dimensional functional network atlas

**DOI:** 10.1101/2021.12.09.472035

**Authors:** Jian Li, Yijun Liu, Jessica L. Wisnowski, Richard M. Leahy

**Author notes:** Co-senior authors (Jessica L. Wisnowski, Richard M. Leahy). Corresponding Author. (R. M. Leahy).

## Abstract

The human brain is a complex, integrative and segregative network that exhibits dynamic fluctuations in activity across space and time. A canonical set of large-scale networks has been historically identified from resting-state fMRI (rs-fMRI), including the default mode, visual, somatomotor, salience, attention, and executive control. However, the methods used in identification of these networks have relied on assumptions that may inadvertently constrain their properties and consequently our understanding of the human connectome. Here we define a brain “network” as a functional component that jointly describes its spatial distribution and temporal dynamics, where neither domain suffers from unrealistic constraints. Using our recently developed BrainSync algorithm and the Nadam-Accelerated SCAlable and Robust (NASCAR) tensor decomposition, we identified twenty-three brain networks using rs-fMRI data from a large group of healthy subjects acquired by the Human Connectome Project. These networks are spatially overlapped, temporally correlated, and highly reproducible across two independent groups and sessions. We show that these networks can be clustered into six distinct functional categories and naturally form a representative functional network atlas for a healthy population. Using this atlas, we demonstrate that individuals with attention-deficit/hyperactivity disorder display disproportionate brain activity increases, relative to neurotypical subjects, in visual, auditory, and somatomotor networks concurrent with decreases in the default mode and higher-order cognitive networks. Thus, this work not only yields a highly reproducible set of spatiotemporally overlapped functional brain networks, but also provides convergent evidence that individual differences in these networks can be used to explain individual differences in neurocognitive functioning.

## Introduction

The human brain is a complex, integrative and segregative network that exhibits dynamic fluctuations in activity across space and time, even at rest. Characterizing this spatiotemporal architecture and the manner in which variations across the population explain individual differences in neurocognitive functions is a central question in neuroscience.

More than two decades ago, scientists first observed that, at rest, the brain exhibits coherent fluctuations in blood oxygenation level dependent (BOLD) signaling – measured by functional MRI (fMRI) – across distributed regions in the human brain (1, 2). From these coherent fluctuations, a canonical set of large-scale resting-state networks have emerged, including the default mode, visual, somatomotor, salience, attention, and executive control networks (3). Evidence suggests that neurocognitive functions emerge from functional interactions within and between these networks and that neurocognitive dysfunction can be, at least partially, understood as an imbalance of activity within and across these networks (4). Although multiple networks are identifiable and highly reproducible across task and resting-state fMRI (rs-fMRI) studies, the computational methods used in their identification have frequently relied on assumptions of orthogonality or statistical independence (5) that are driven more by their appealing mathematical properties than our understanding of brain networks. As a result, we may inadvertently constrain the characteristics of identified networks and consequently, our understanding of the human connectome.

Two distinct approaches to identification of brain networks are described in the literature: (i) those based on spatial decompositions in which each network is represented as an image reflecting the participation of each voxel (or surface element in cortical-only maps); (ii) those based on parcellation in which the brain (or more frequently just the cerebral cortex) is partitioned into a series of non-overlapping regions that represent functionally homogeneous regions within the brain. In the former group, independent component analysis (ICA) is most often used for brain network identification. Among many variations, spatial ICA is the most frequently used approach for multi-subject fMRI data. ICA is applied to temporally concatenated data from individuals with an independence constraint in the spatial domain to identify common networks across subjects (5). Many meaningful components can be extracted using these ICA-based approaches (6). Although these methods allow temporal correlations among networks, the estimated spatial maps are statistically independent. An alternative approach uses seeded correlation to identify regions in the brain whose resting brain activity is correlated with the seed location (1, 7). While networks identified using different seed locations do not result in the independence property observed with ICA, seeded correlation does not offer a systematic approach to identification of a complete set of networks since the networks found are dependent on the selection of seed locations.

As an alternative approach, Yeo *et al*. parcellated the human cortex based on the Pearson correlation of rs-fMRI time-series between cortical vertices (8). Follow-up studies extended the parcellation framework to whole-brain (9), a larger number of parcels (10), or multi-modal datasets (11). Others explored similarity measures other than the Pearson correlation, including homogeneity and boundary maps (12, 13). The parcellation approach emphasizes functional homogeneity within each parcel and explicitly prohibits spatial overlap. Therefore, a contiguous cortical region that is fully involved in one network but only a portion of which is involved a second, needs to be represented by two parcels. This leads to the need for very dense parcellations to accommodate overlapping and interacting networks. An alternative to identifying spatially homogeneous regions is to use a sliding-window framework for identification of quasi-stationary brain macro-states. Methods have been described using *k*-means clustering (14), graph analysis (15), and dictionary learning (16). These sliding-window-based methods assume the brain is in a single “state” at each time point, so in contrast to the spatial parcellation methods, these methods instead perform a non-overlapping parcellation in time.

Functional connectivity has been shown to fluctuate over time in a manner that is more complex than the discrete macro-state model (15, 17–19). Assumptions of stationarity in time may therefore be too restrictive to capture the dynamic nature of the brain (20). On the other hand, spatial overlap between large-scale brain networks has been widely observed in fMRI data (18, 21). Supporting evidence shows that cortical brain activities are not segregated in space, rather they overlap and interact with each other at different scales, which could potentially be attributed to the heterogeneous activity of intermingled neurons within the same brain region (22). Specifically, the laminar organization of the cerebral cortex combines a hierarchical structure with a high degree of parallel processing (23). The extrinsic connections to and from the cerebral cortex target specific layers within the supragranular layer (layers 2/3), granular layer (4), and infragranular layers (5/6) depending on the loci of origin and termination, and the hierarchical relationship between the regions. Furthermore, neural electrophysiological oscillations (alpha, beta, and gamma-band) appear to have a laminar pattern, with gamma-band activity predominantly observed in superficial layers while alpha and beta-band activity is stronger in deep layers. (24–27) Together, this microstructural organization suggests that at a macroscale, cortical and subcortical regions should interact across multiple networks superimposed across space and time. However, to date, no study has defined a taxonomy of functional brain networks that fully incorporate the spatiotemporal properties of rs-fMRI signals.

In this work, we define a brain “network” as a functional component that jointly describes the spatial distribution and temporal dynamics, where neither the spatial nor the temporal domain is unrealistically constrained. In contrast to the traditional network terminology, e.g., the default mode network, where only the spatial map is well characterized, this renewed definition could help us explore one fundamental question in functional connectivity analysis: which parts of the brain (the question of “where”) talk to each other at which given period of time (the question of “when”). We approach this network identification problem by first temporally synchronizing rs-fMRI data to a common space via the BrainSync algorithm (28, 29) because rs-fMRI time series are not directly comparable across subjects. BrainSync exploits correlation similarity across subjects to perform an orthogonal rotation of rs-fMRI time series, such that the data in homologous regions of the brain are highly correlated after synchronization. Synchronized group rs-fMRI data are then combined using an additional subject dimension to form a third-order tensor (space × time × subject). Brain networks are modeled as low-rank Canonical Polyadic components and identified using our recently developed Nadam-Accelerated SCAlable and Robust (NASCAR) tensor decomposition method (30, 31).

We identified twenty-three brain networks using the rs-fMRI data from a large group of healthy subjects acquired by the Human Connectome Project (HCP) (32, 33). These networks are spatially overlapped, temporally correlated, and highly reproducible across two independent groups and sessions. Based on coherence patterns in the spatial, temporal, spectral, and subject modes, we show that these networks can be clustered into six distinct functional categories. One category contains global signals with four different spatial patterns, which are related to physiological rather than the “cognitive” components. The remaining five categories included networks that have been previously associated with the default mode, visual, somatomotor, auditory, and higher-order cognitive functions. The spatiotemporal organization of the brain networks identified by the NASCAR method naturally formed a representative functional network atlas for the healthy population. Furthermore, NASCAR also provides a “subject mode” network participation level for each subject from the third dimension of the tensor. To determine whether individual differences in these networks, represented in the subject mode, can explain differences in cognitive function, we projected an independent attention deficit hyperactivity disorder (ADHD) rs-fMRI dataset from Peking University as part of the ADHD-200 dataset (34) onto the network atlas formed from the spatial and temporal components of each network identified from the HCP data. Using the resulting subject mode, representing the level of activity for each subject in each network, we were able to differentiate ADHD from neurotypical (NT) subjects. The classification accuracy outperformed state-of-the-art methods by a large margin. Furthermore, we demonstrated that individuals with ADHD displayed disproportionate brain activity increases, relative to NT, in visual, auditory, and somatomotor networks concurrent with decreases in the DMN and higher-order cognitive networks. Taken together, this work not only yields a highly reproducible set of spatiotemporally overlapped functional brain networks (the “network atlas”), but also provides convergent evidence that individual differences in these networks can be used to explain individual differences in neurocognitive functioning.

## Results

### Highly reproducible brain networks were identified from resting-state fMRI data

We analyzed the minimally preprocessed 3T HCP rs-fMRI dataset from 1000 healthy subjects (32, 33). The rs-fMRI data were first temporally synchronized across subjects using the BrainSync algorithm (28, 29). This orthogonal transform results in highly correlated time-series at homologous locations across subjects. We then applied NASCAR tensor decomposition (30, 31) to the synchronized rs-fMRI data to identify a set of distinct brain networks, each consisting of a spatial map, time series, and subject mode (or participation level). The subjects were randomly assigned into two groups, 500 subjects each, with two 15 min sessions per subject. NASCAR was applied separately for each session of the two groups to produce a total of four separate network decompositions. Identical parameters were used in each case with a maximum of 50 components. The resulting NASCAR components from the other three groups/sessions were best matched to that from the first session of the first group using the Gale-Shapley algorithm (35). Through visual examination of the NASCAR components in the first group first session, seventeen subject-specific or physiologically implausible “noise” components at higher ranks were removed from further consideration. Among the remaining networks, we identified twenty-three spatially overlapped, and temporally correlated brain networks that are highly reproducible across the two large independent groups as well as the two independent sessions. Inclusion in this group required a maximum inter-session/inter-group Pearson correlation greater than 0.9 across the network’s spatial map as shown in Fig. S1 in the Supporting Information. These twenty-three networks were then categorized into six functional categories based on visual examination of their cross-correlations in space, time, spectrum, and subject mode, where the spectrum was estimated for each network using the Welch method (36). Figs. 1 and 2 show these twenty-three networks grouped by their functional categories enclosed in different color boxes. Figs. S2 and S3 in the Supporting Information show the same set of networks with inflated surface representations. See Fig. 3 below for their correlation matrices and the Materials and Methods section for the detailed processing pipeline.

**Fig. 1.**
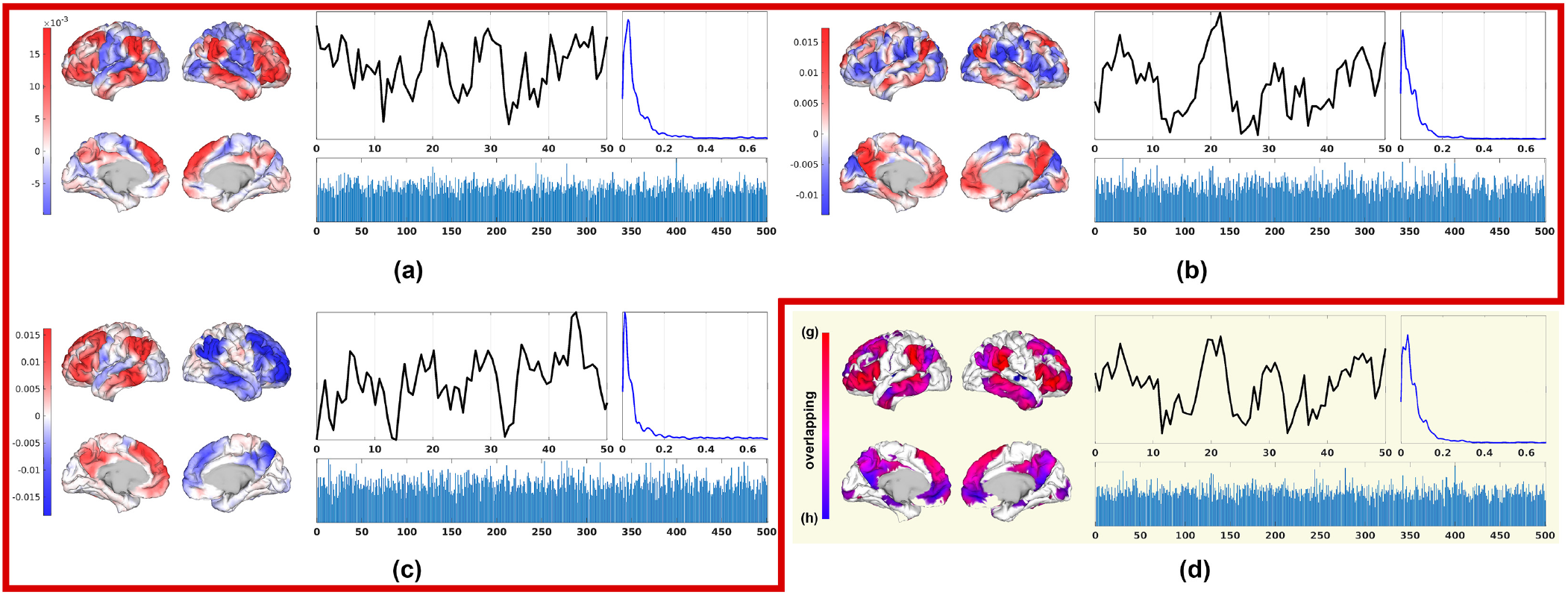
Subset of the twenty-three identified brain networks that belongs to the default mode category. Red (a) – (c): Three sub-networks of the default mode network; (d) The combined network from (a) and (b), where (a) is encoded in red, (b) is encoded in blue, and purple indicates overlapped regions. In each network, the spatial map is shown on the left with a bipolar color scheme where red represents coherent and blue anti-coherent regions; temporal dynamics (in secs) are shown on the top middle, where the time series is truncated to 50 seconds for visual clarity; the subject participation level is shown on the bottom and Welch-estimated power spectra (in Hz) of the temporal mode are shown top-right. The peak frequencies in the spectrum are listed in Table S1 in the Supporting Information.

**Fig. 2.**
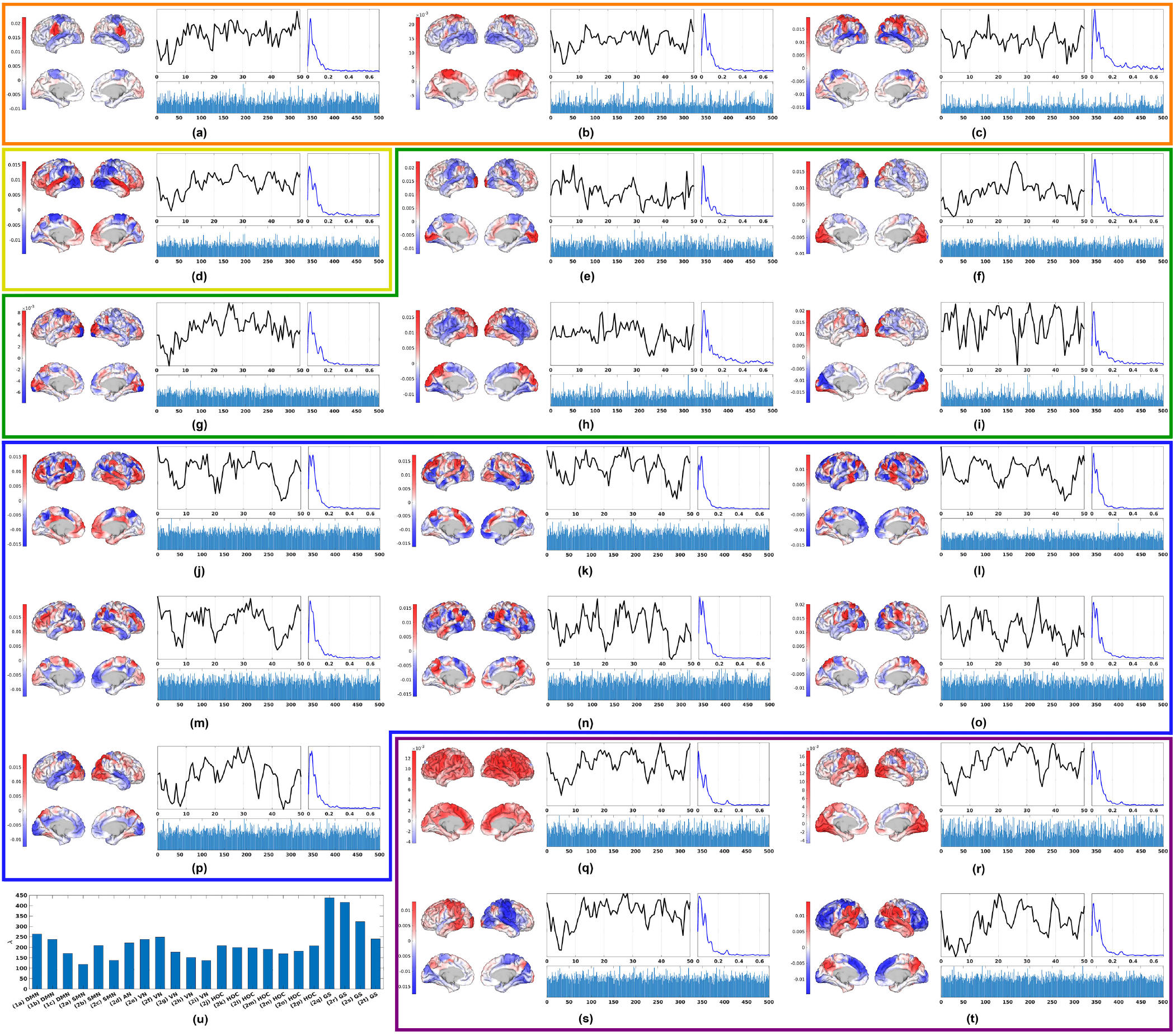
Twenty of the twenty-three identified brain networks grouped by their functional categories. Orange (a) – (c): somatomoter networks; Yellow (d): auditory network; Green (e) – (i): visual networks; Blue (j) – (p): higher-order cognitive networks; Purple (q) – (t): global signals. See Fig. 1 for the three DMN networks and description of components shown for each network. (u) Plot of the *λ*values for each network in Figs. 1 and 2 representing the relative degree of activity in each network averaged across the population. Individual strength of participation in each network is found by scaling of *λ*by the subject participation level.

**Fig. 3.**
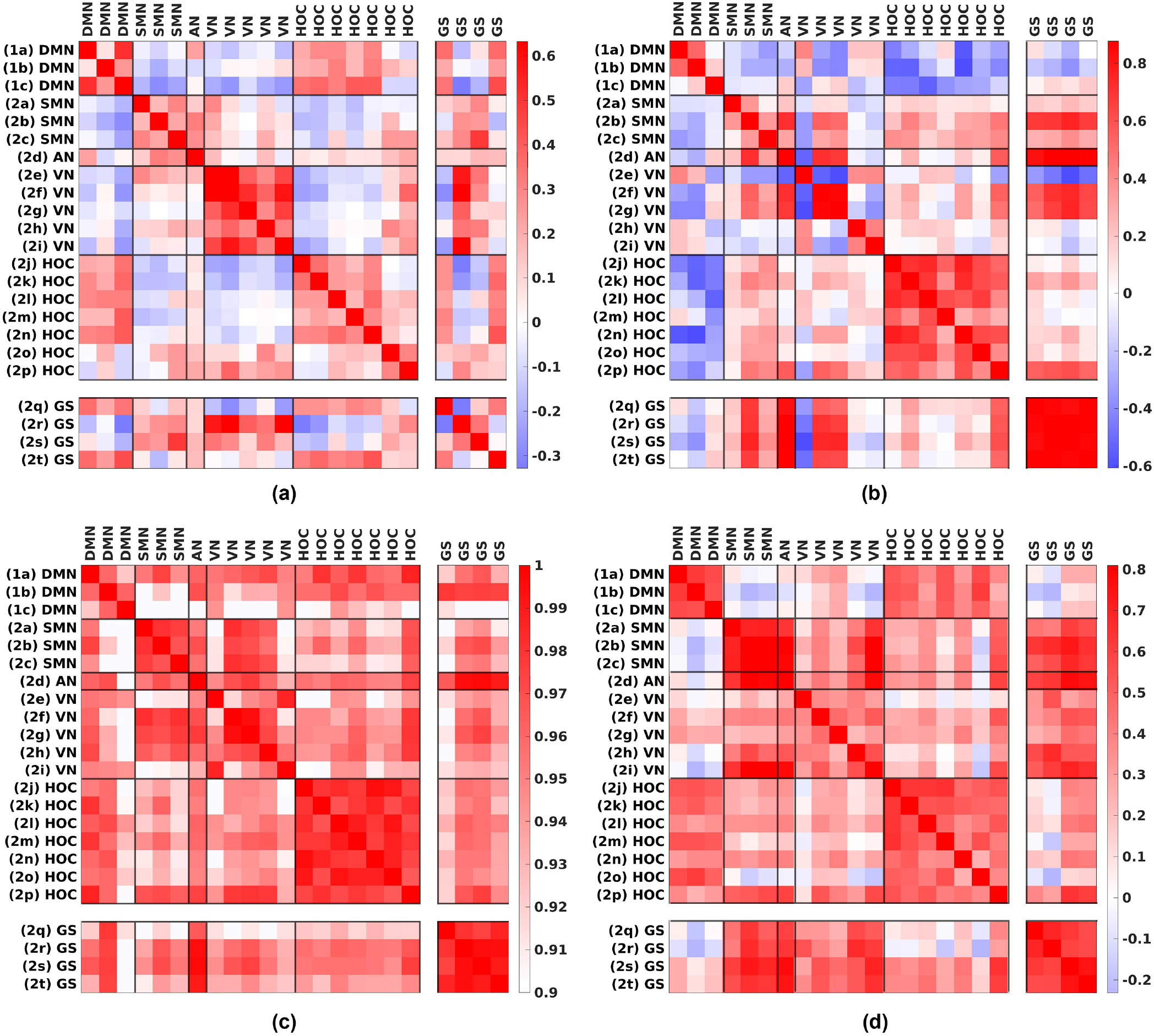
Cross-correlation matrices of the twenty-three networks/components: (a) spatial correlation; (b) temporal correlation; (c) spectral correlation; (d) subject correlation. The labels on the axes indicate the functional category each network belongs to: DMN – default mode network; SMN – somatomotor network; AN – auditory network; VN – visual network; HOC – higher-order cognitive network; GS – global signal. The indices inside the parentheses correspond to the sub-figure indices in Figs. 1 and 2. The color bar on the right shows the range of correlation values in each individual matrix.

Three networks, Fig. 1 (a) – (c), included the core regions of the canonical DMN: the medial prefrontal cortex (mPFC), precuneus, posterior cingulate cortex (PCC), the inferior parietal lobule (IPL), and the lateral temporal cortex (LTC) (superior temporal sulcus and middle temporal gyrus). Spatially, these networks displayed distinct features: Network (a) showed the strongest coherence between the dorsomedial (dm)-PFC, rostral IPL (supramarginal gyrus), LTC, and the inferior frontal gyrus (IFG). In contrast, network (b) showed the strongest coherence between the medial PFC (ventral to the dmPFC in (a)), precuneus/PCC, and caudal temporoparietal junction (TPJ) (angular gyrus). Network (c) showed strong spatial overlap with both (a) and (b) but with an opposite sign between the left and the right hemispheres, indicating an oscillatory left-right pattern. All three networks had positive temporal correlations with each other and showed strong correlation in the subject mode, Fig. 3 (d). We also note that these networks appear to operate at different frequencies as shown in their spectrum: (a) has a higher peak frequency (0.027 Hz) relative to (b) and (c) (0.011 Hz), Table S1. Overall, the patterns revealed in Figs. 1 and 3 suggest that (a) and (b) may represent sub-networks of the canonical DMN. To test this hypothesis, we combined (a) and (b) via a weighted sum by their network strengths (*λ* value in Eq. 1). As shown in Fig. 1 (d), the merged network strongly resembles the canonical DMN as obtained using seeded correlation or ICA-based approaches.

Three networks, Fig. 2 (a) – (c), included core regions of the somatomotor cortices. Networks (a) and (b) centered on the face/tongue and foot areas, respectively, while network (c) was centered on the hand region, premotor, supplementary motor, and sensory cortices. These somatomotor networks (SMN) show modest correlations in the spatial and temporal mode, Fig. 3 (a) and (b), but strong coherence in their spectrum with a common peak frequency at 0.027 Hz, Fig. 3 (c) and Table S1. Across subjects, the large discrepancy suggests that there was a subset of participants who were either more active physically during the rs-fMRI sessions or more aware of their bodies and actively focused on keeping still. However, this cross-subject discrepancy was very consistent among the three networks, as reflected by the subject mode correlation in Fig. 3 (d).

One network, Fig. 2 (d), was centered on primary auditory cortices (AN), including Heschl’s gyrus and the surrounding regions of the superior temporal gyrus. This network was distinct from the remaining 22 networks in the spatial mode, however, the AN is correlated with multiple other networks in the temporal, spectral, and subject mode (Fig. 3).

Five networks, Fig. 2 (e) – (i), were centered on visual cortices (VN). Network (e) was centered on the primary visual cortices (striate cortices) along the calcarine fissue (V1, Brodmann’s area [BA] 17) while network (f) included V1 as well as secondary and tertiary areas (extrastriate cortices) in the cuneas and lingual gyri (V2, V3 and VP, BA 18 and 19). Network (g) included secondary and tertiary areas in cuneus, lingual and lateral occipital gyri as well as the TPJ. Networks (h) and (i) were centered in the visual association areas and showed a strong correspondence, spatially, to the dorsal and ventral visual pathways, respectively. Temporally and spectrally, Fig. 3 (b), (c) and Table S1, these five networks appeared to segregate further into three smaller clusters (e, f/g, h/i) suggesting that spatiotemporal interactions across the primary, secondary/tertiary, and visual association networks differ. Modest correlations were observed across all five networks in the subject mode, Fig. 3 (d).

Seven networks (Fig. 2 (j) – (p)) were centered on higher-order association cortices in the frontal, temporal, and parietal lobes (HOC). Networks (j) and (m) had nodes in the lateral prefrontal cortex, inferior parietal lobule (supramarginal gyrus and/or intraparietal sulcus), and inferior temporal lobe, which are well aligned with the central executive control network. Core regions of network (k), (l), and (n) included anterior midcingulate cortex, bilateral anterior insula, inferior parietal lobule, temporal parietal junction. These three networks correspond to the cingulo-opercular, salience, and ventral attentional network, respectively and may collectively represent the large-scale midcingulo-insular system (3). Networks (o) and (p) were centered on the superior parietal lobule extending posteriorly along the intraparietal sulcus into lateral occipital cortex, including areas V3A and the middle temporal complex (MT+), while network (o) also includes prefrontal regions in the vicinity of the frontal eye fields and inferior frontal junction. Together these two networks are close to the canonical dorsal attentional network. In fact, these two networks were spatially segregated from the other HOC networks as shown in Fig. 3 (a). However, all HOC networks were highly correlated in their temporal, spectral, and subject modes, Fig. 3 (b) – (d). In particular, they showed strong consistency in bi-modal operating frequencies with the lower at 0.011/0.016 Hz and the higher at ∼0.04 Hz, Table S1.

In contrast to the above, the final four “networks”, Fig. 2 (q) – (t), appear to collectively represent the global signals (GS) present in the rs-fMRI data (the HCP minimal preprocessing pipeline does not include a global signal regression step). The Pearson correlations between the four components and the empirically computed global signal are 0.98 for (q), 0.90 for (r), 0.81 for (s), and 0.87 for (t), respectively. The component (r) has the highest activity level (*λ*) among all networks, including the DMN, Fig. 2 (u). They each exhibit variable spatial patterns with emphasis on different regions of the brain, but (q) has a more global distribution than the other three. Temporally and spectrally, we observed a very high concordance among the four components, Fig. 3 (b) and (c), where they all have three distinct peaks, Table S1. The first peak frequency of (q) was lower (0.005 Hz) than the other three (0.016 Hz), the second peak was around 0.071/0.076 Hz, and interestingly, they all showed a consistent unique third peak at 0.288 Hz, which is substantially higher than peak frequencies of all other networks and corresponded to a respiration rate of 0.288 Hz, approximately equivalent to 17 breaths per minute (this particular frequency is specific to the subject who was used to synchronize the time series back to in order to estimate its spectrum, see the Method section). They were also consistent in the subject mode, Fig. 3 (d).

### Intrinsic brain networks are spatially overlapped and temporally correlated

The interactions and relations among networks persist in multi-dimensional facets, including space, time, spectrum, and the subject participation level. Moreover, the inter-network relationship spans a hierarchical organizing structure: Networks within a classical functional category (intra-category) have their unique means of interaction as discussed in the previous section and shown in the six diagonal blocks of Fig. 3. At the same time, networks in one functional category have consistent but differing relations, also in a multi-dimensional manner, to networks in other categories (inter-category) as seen in the off-diagonal blocks in Fig. 3. Of the many inter-categorical relations, the most evident one is the strong and consistent temporal anti-correlations between the DMN and the HOC, Fig. 3 (b). This anti-correlation between “task-positive” networks and the DMN, which is also called “task-negative” network historically, has been widely reported in the literature (37–40). Further, the spatial distribution of networks involved in early developed regions, such as SMN, VN, AN, tend to be anti-correlated to that involved in later developed HOC and DMN, as shown in Fig. 3 (a). Similar segregation can be observed in the subject mode, where subjects who have higher participation level in the DMN also have higher participation level in HOC, but lower in SMN, AN, and VN, while interestingly the GS components have consistent behavior with the latter, Fig. 3 (d). Last but not the least, regardless of the positive and negative correlations in space, time and the subject mode, networks more or less were operating in similar frequencies, with higher consistency within each functional category and lower between categories, Fig. 3 (c) and Table S1.

To better illustrate and visualize how these spatially overlapped and temporally correlated networks interact with each other, we mapped the spatiotemporal patterns (the outer product of the spatial map and the temporal dynamics) of the two sub-networks of the DMN, Fig. 1 (a) and (b), onto a common tessellated surface. They were then color-coded and played back in real time as a video shown in Movie S1 in the Supporting Information (This is similar to how we combined the two sub-networks of the DMN in Fig. 1 (d) but now in a time evolving manner).

As a comparison, we ran the commonly used group ICA algorithm (5) on the same dataset and obtained fifty common spatial maps. The temporal dynamics were then back-reconstructed per (5, 6), and the spectra were estimated using the Welch method with identical parameters to those used in the spectrum estimation for NASCAR. We then similarly computed the spatial, temporal, and spectral cross-correlation matrices as shown in Fig. S4 in the Supporting Information. The network order was matched to the NASCAR results using the Gale-Shapley algorithm. The correlation matrix, Fig. S4 (a), between the spatial maps is an identity matrix, which is expected as spatial ICA imposes independence constraints on the spatial maps. Clusters of networks are present in the temporal and spectral correlation matrices, Fig. S4 (b) and (c), but the patterns are less clear than those in the NASCAR results. Further, as with other 2D methods, group ICA does not directly produce the third dimension capturing the subject participation level, which is present in the 3D NASCAR model as shown in Fig. 3 (d) (although a similar estimate could be extracted by computing a two norm on the time series for each network and subject).

Fig. S5 shows the cumulative variance explained of PCA, NASCAR, and ICA. The components of PCA, as expected, provide an upper bound on the variance explained because of the orthogonality of spatial and temporal modes between networks and the property of the singular value decomposition used to compute the PCA that it maximizes variance explained across all possible decompositions at a given rank (41). However, such an orthogonality condition is unlikely to be biologically plausible for brain networks, and thus NASCAR does not enforce such constraints. Compared with components from spatial ICA, NASCAR consistently has a higher cumulative variance explained, even though it has shared modes in both space and time (after BrainSync transformation), unlike spatial ICA from which we compute subject-specific temporal modes using least squares.

### The spatiotemporal networks build a functional network atlas

In contrast to brain atlases where the brain is parcellated into many spatially contiguous, non-overlapping, and hopefully functionally distinct and homogenous regions, the networks identified by the NASCAR method above naturally form a spatiotemporal atlas that jointly describes overlapped spatial maps and correlated temporal dynamics, which we refer to as a “functional network atlas”.

Here we illustrate one of many potential applications of this atlas via an rs-fMRI study of ADHD subjects. We used independent rs-fMRI data acquired from Peking University as part of the ADHD-200 dataset (34, 42), where 95 subjects are classified as ADHD and 126 are neurotypical (NT) subjects as the control group. The preprocessed rs-fMRI data were synchronized and projected onto the atlas and the subject participation level for each network was obtained using a least-square fit. The ADHD diagnostic label (binary) and index (continuous-valued) were also provided by (34), where the ADHD index was measured by the ADHD Rating Scale IV, a combined measure of inattention and hyperactivity-impulsivity. Because of intrinsic variance in the ADHD index measures and low SNR in individual rs-fMRI data, prior to analysis eleven outliers were identified and removed based on Cook’s distance metric in a one-pass linear regression model using the estimated subject participation level and the ADHD index (see Materials and Methods section for detail). The estimated subject participation levels (one per network per subject) embed the intrinsic properties of the brain networks for each subject. The concatenation of participation levels from all subjects forms a feature matrix ***C*** ∈ ℝ^*N*×*R*^ as shown in Fig. 4 (a), where *N* is the number of subjects and *R* is the number of networks. To demonstrate the effectiveness of information extraction from the rs-fMRI data and embedding in this parsimonious feature matrix, we show below how this feature matrix ***C*** could be used to 1) predict ADHD index; 2) classify ADHD subjects from NT subjects; and 3) provide insight into the differences in network properties between ADHD and NT.

**Fig. 4.**
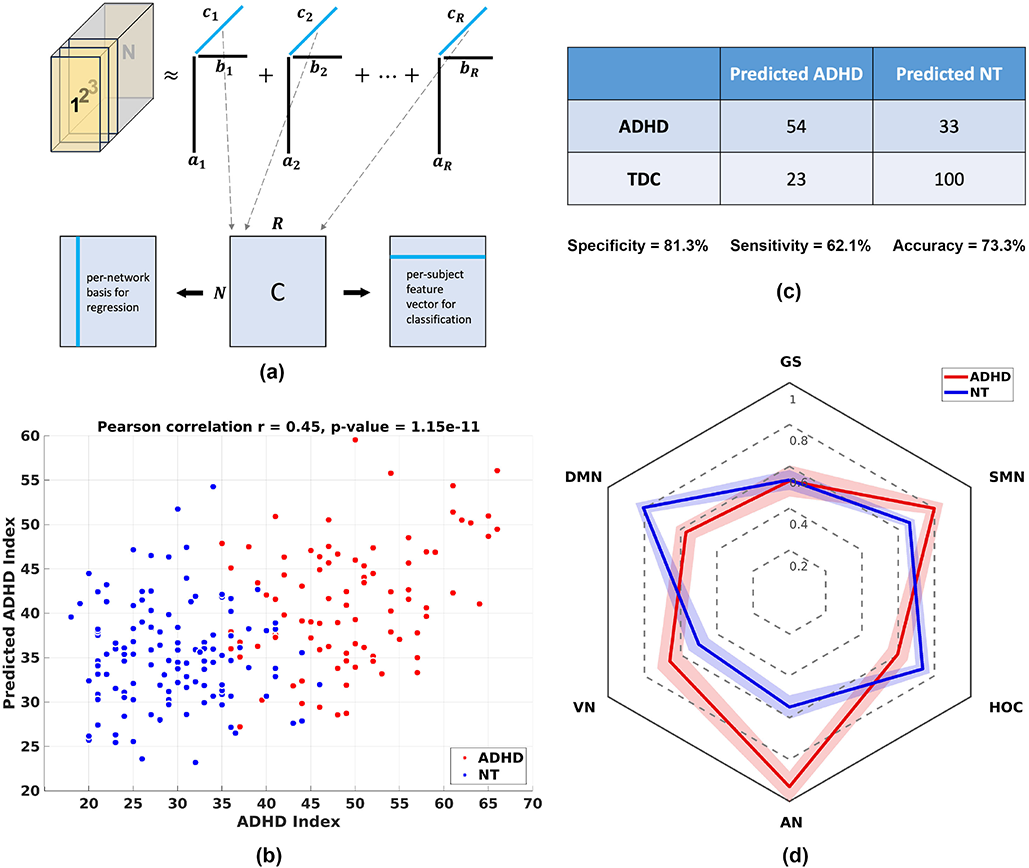
Results of applications of functional network atlas to the ADHD dataset. (a) The feature matrix ***C*** is formed by concatenation of the estimated subject participation scores. It could be used as per-network bases for regression study (left arrow) or per-subject feature vectors for classification tasks (right arrow); (b) Scatter plot between the predicted ADHD index (y-axis) and the measured ADHD index (x-axis). Red dots mark the ADHD subjects, and the blue dots mark the neurotypical (NT) subjects; (c) The binary confusion matrix of the classification result, with accuracy, specificity, and sensitivity reported below the table; (d) The radar plot of the (normalized) median subject participation level in six groups of networks for ADHD in red and NT in blue. The shaded area under the hexagons represents their corresponding standard errors.

We first treated the feature matrix ***C*** as a group of column vectors, one per network (Fig. 4 (a) left arrow). We then used these column vectors as bases to predict the ADHD index using linear regression with 10-fold cross-validation. Fig. 4 (b) shows the scatter plot between the ADHD index (y-axis) and the measured ADHD index (x-axis) predicted from the held-out (cross-validation) data, where the ADHD and NT subjects are colored in red and blue respectively. The Pearson correlation between these two measures is 0.45 with a *p*-value of 1.15 ×10^−11^. To our knowledge, most fMRI-based ADHD studies focus on either the classification problem or studies of differences in brain networks in ADHD subjects relative to NT subjects, as discussed below. We are unaware of any previous study that predicts ADHD index, except for our previously reported correlation of 0.32 (43), which is based on a deep neural network

We next used the row vectors in feature matrix ***C*** (Fig. 4 (a) right arrow) as features to differentiate ADHD from NT. Specifically, we trained a binary support vector machine (SVM) (44) with a linear kernel using the ground truth diagnostic/label (we do not consider sub-types of ADHD here) provided by the ADHD-200 Consortium to classify ADHD and NT. Fig. 4 (c) shows the 10-fold cross-validated classification result as a binary confusion matrix. We are able to achieve 73.3% classification accuracy with a balanced 81.3% specificity and 62.1% sensitivity. In contrast, using the same Peking University dataset, the best performance obtained in ADHD-200 global competition across all teams was 51.1% (42). It also substantially outperforms more recent convolutional neural network-based studies using rs-fMRI data only (45, 46), using a combination of rs-fMRI and structural scans (47), where the highest reported accuracy was 65.2% (46).

To illustrate how activity in these networks differs between ADHD and NT, we took the average subject participation level in each of the six network categories (GS, DMN, VN, AN, HOC, and SMN) for ADHD and NT separately and computed their medians and standard errors. The results shown in Fig. 4 (d) were normalized identically between the ADHD and NT to have a maximum of 1 and a minimum of 0.5 and displayed using a radar plot for easy visualization and comparison (48). Consistent with the literature, ADHD subjects present a decreased activity level compared to NT in DMN (49) as well as networks related to cognitive functions (50). On the other hand, VN, AN, and SMN show an increased level of activity (51) in ADHD subjects relative to NT. This is as we expected as ADHD subjects tend to be more inattentive and physically hyperactive than NT. We do not see a significant difference in the global components between the two.

## Discussion

The present study deepens our understanding of the intrinsic spatiotemporal organization of the human connectome by describing, for the first time, a functional atlas consisting of a set of twenty-three resting-state networks with associated spatial and temporal maps. Individual subjects’ fMRI can be mapped to this atlas by using BrainSync to synchronize their time series to that of the atlas, followed by a least-squares fit to the spatial-temporal model formed by the 23 networks. Critically, these networks were derived without the mathematical constraints imposed in previous studies, while demonstrating high reproducibility across two large, independent groups of subjects as well as between two independent sessions in each group. Furthermore, these networks present nearly symmetric and spatially contiguous spatial patterns. This is particularly encouraging because, unlike 2D (space × time) methods such as ICA (5), there is no spatial constraint imposed in the low-rank tensor model (30, 31). We also note that we did not apply any additional spatial smoothing beyond the 2 mm isotropic Gaussian smoothing performed by the HCP minimal preprocessing pipeline (32). Hence, the symmetry and spatial contiguity of the networks very likely reflects an intrinsic property of these human brain networks. In addition, the spatial topography of these multi-dimensional networks resembled many of the canonical resting-state networks (RSNs), including the DMN (37–40), the visual networks, and the higher-order cognitive networks (3).

A unique property of the tensor-based NASCAR method is that it intrinsically includes an extra (third) dimension that encodes rs-fMRI information for each subject. In contrast, group-ICA and related methods typically concatenate the time series at each location for all subjects into a single dimension to create a second-order tensor (or matrix). While it is possible to compute a per-subject participation value once ICA networks are found (through computing the relative power in the time series for a given network for each subject), these methods do not exploit any *intrinsic* low-rank structure embedded directly in the third-order tensor. An analogy to this situation is image compression using JPEG: high compression rates using 2D cosine transforms are achieved by exploiting intrinsic spatial structure in (2D) images. If the images were instead concatenated into a single 1D vector, far lower compression rates would be realized using the corresponding 1D transform). In fact, it is this embedding into a third-order rather than a second-order tensor to exploit low-rank structure that allows relaxation of the independence assumption required to obtain meaningful decompositions in the second-order case. As a result, the ability to consistently find networks that exhibit both spatial and temporal correlations is intrinsic to the use here of the third-order tensor decomposition. To arrange the rs-fMRI in this way requires similarity in time-series across subjects. While each subject exhibits independent resting activity in the rs-fMRI scan, applying the BrainSync transform prior to tensor decomposition exploits similarity in correlation across subjects in order to align the time-series across individuals at homologous locations.

### Spatiotemporal organization of the default mode network and subsystems

We observed three multi-dimensional RSNs related to the DMN (Fig. 1). Two of these, Fig. 1 (a) and (b), were bilaterally symmetric, and when combined by weighting by their relative strengths (*λ*in Eq. 1) resembled the canonical DMN, as shown in Fig. 1 (d), suggesting that individually they represent subsystems of the DMN. Prior studies have demonstrated that the DMN can be decomposed into at least two partially-distinct subsystems that are inter-connected to and interact with each other (37, 38). Using a hierarchical clustering approach, these have been described as: a “dorsal medial subsystem” which consists of the temporal pole, the lateral temporal cortex, the temporal-parietal junction, and the dorsomedial prefrontal cortex and a “medial temporal subsystem” which consists of the hippocampal formation (HF), the parahippocampal cortex, the retrosplenial cortex, the posterior inferior parietal lobule (pIPL), and the ventral medial prefrontal cortex (PFC) (7, 38, 52). The posterior cingulate cortex (PCC) and the anterior medial prefrontal cortex have been strongly associated with both subsystems. The spatial topography of Fig. 1 (a) and (b) strongly resemble the dorsal and medial temporal subsystems, respectively. Furthermore, our results show that the dorsal and medial temporal subsystems have different spectral characteristics, Table S1, with the dorsal medial subsystem demonstrating peak power at a higher frequency as compared to the medial temporal subsystem. A prior study, utilizing repetitive transcranial magnetic stimulation (rTMS) and fMRI, demonstrated that two different frequencies of rTMS applied to the same DMN node in the pIPL induced two topographically distinct changes in functional connectivity (53). High-frequency rTMS to the pIPL decreased functional connectivity between the PCC and medial PFC (mPFC) but not between those nodes and the HF. In contrast, low-frequency rTMS to pIPL did not alter connectivity between the PCC and mPFC, but did increase connectivity between the pIPL and the HF. Rather than a single node possessing multiple, functionally distinct relationships among its distributed partners, our results suggest that a single node may belong to two or more spatiotemporal networks that are superimposed on top of each other. Across time (Movie S1, Supporting Information), the different frequencies result in what appears to be a cyclical pattern – alternating between periods of co-activation and periods where they are individually activated; however, this is likely epiphenomenal – an effect of multiplexing.

In addition to the two networks described above, our results revealed a third default mode-related network. The spatial topography of this network was nearly identical to the above; however, spatiotemporally, there were strong correlations across the hubs within hemisphere and anti-correlations between hemispheres. This spatiotemporal pattern has not, to our knowledge, been reported in other rs-fMRI studies; however, it is consistent with the temporal dynamics of the DMN as demonstrated by magnetoencephalography (MEG) (54). Similar to our findings, that study demonstrated non-stationarity of the MEG activity with transient formation of complete DMN, suggesting that the stationarity of RSNs demonstrated in prior rs-fMRI studies may be more directly related to the methods employed rather than the underlying physiology of the RSNs. Overall, the findings from this study support that the canonical DMN observed in prior rs-fMRI studies emerges statistically from the spatiotemporal interactions across multiple DMN subsystems. These new observations about the spatiotemporal relationship between the two sub-networks could potentially provide a deeper insight into the functional organization of the human DMN as well as how alterations in the spatiotemporal dynamics between these subsystems might contribute to cognitive dysfunction (52, 55, 56).

### Somatomotor, auditory, and visual networks

Three networks, Fig. 2 (a) – (c) included core regions of the somatomotor cortices. Networks (a) and (b) centered on the face/tongue and foot areas, respectively, while network (c) was centered on the hand region, premotor, supplementary motor, and sensory cortices. In the canonical 7-network RSN parcellation by Yeo *et al*, the somatomotor network is a single network encompassing motor and somatomotor cortices anterior and posterior to the central sulcus extending into Heschl’s gyrus and the dorsal aspects of the superior temporal gyrus. Additionally, at least two subsystems have been observed: including right, left (Smith et al. 2009), dorsal (hand) and ventral (face) subsystems (8, 12) as well as separation of auditory and somatosensory face areas (Kong et al. 2019). Spatially, our results are similar to the higher-order parcellation schemes, and further demonstrate modest correlations in the spatial and temporal mode, Fig. 3 (a) and (b), but strong coherence in their spectra with a common peak frequency at 0.027 Hz across all somatomotor networks, which, in turn, is readily dissociable from the spectral characteristics of the auditory network (Table S1).

Five networks included core components of the visual system. Three of these networks were centered in the primary, Fig. 2 (e), and secondary and tertiary visual cortices (f, g) in the occipital lobe, while two (h, i) centered in visual association areas and showed a strong correspondence, topologically, to the dorsal and ventral visual pathways, respectively. Prior studies have demonstrated various occipital networks, including medial and lateral networks, as well as one corresponding to the occipital pole. However, to our knowledge, this is the first study to identify RSNs corresponding to the dorsal (“where”) and ventral (“what”) visual pathways first proposed 40+ years ago on the basis of neuropsychological, electrophysiological, behavioral evidence (57, 58), and later, structural connectivity (59). Furthermore, our results demonstrate unique spectral characteristics across the visual networks. First, each of the visual networks exhibited a peak at 0.06 Hz. This peak was unique to the visual networks and was not observed across any other networks. Additionally, the primary and tertiary peaks, appeared to cluster, in line with the spatial topography of the networks across the primary, secondary/tertiary, and visual association cortices. Likewise, the networks corresponding to the primary visual (e) and auditory (d) exhibited peak power at the same frequency (0.016 Hz). Again, these findings suggest that elucidating the spectral characteristics of the RSNs may be key to understanding how information is represented or transmitted across the human brain.

### Higher-order cognitive networks

Seven networks, Fig. 2 (j) – (p), were centered on higher-order association cortices in the frontal, temporal, and parietal lobes. Given this spatial topography, we refer to these as the Higher-Order Cognitive (HOC) networks. Networks (j) and (m) had nodes in the lateral prefrontal cortex, inferior parietal lobule (supramarginal gyrus and/or intraparietal sulcus), and inferior temporal lobe, which are well aligned with the central executive control network (3). In contrast, networks (k), (l), and (n) included anterior midcingulate cortex, bilateral anterior insula, inferior parietal lobule, temporal-parietal junction. These three networks corresponded to the cingulo-opercular, salience, and ventral attentional network, respectively, and may collectively represent the large-scale midcingulo-insular system (3). Networks (o) and (p) were centered on the superior parietal lobule extending posteriorly along the intraparietal sulcus into lateral occipital cortex, including areas V3A and the middle temporal complex (MT+), while network (o) also includes prefrontal regions in the vicinity of the frontal eye fields and inferior frontal junction. Together these two networks are close to the canonical dorsal attentional network. In fact, these two networks were spatially segregated from the other HOC networks as shown in Fig. 3 (a). However, all HOC networks were highly correlated in their temporal, spectral, and subject modes, Fig. 3 (b) – (d). In particular, they showed strong consistency in their bi-modal operating frequency, with the lower one at 0.011/0.016 Hz and the higher one at ∼0.04 Hz, Table S1. These spectral characteristics suggest that while the HOC networks represent different subsystems, they operate at, and therefore may interact, at specific frequencies. Furthermore, while the higher frequency (∼0.04 Hz) is unique to the HOC, the lower frequencies were shared with other networks, including subsystems within the DMN, auditory and visual networks, suggesting a mechanism by which the attentional and executive subsystems may influence information processing across other subsystems.

### Global signals

The four global components, Fig. 2 (q, r, s, t), exhibit strong correlations with the global signal computed from the rs-fMRI data empirically, which is often used in global signal regression. Indeed, all four networks exhibit spectral peaks at 0.28Hz, which is consistent with an average respiration rate of 17 per min. These global signals can be attributed in part to vascular effects or neurovascular coupling caused by respiratory and/or cardiac activity (60–62). In particular, Fig. 2 (r) shows a very similar spatial map to the “physiological network” reported in (63), which is obtained directly by a linear regression using physiological recordings. In contrast, NASCAR identified these four global signals in a purely data-driven manner without access to physiological measures. Further, it has been shown that physiological components cannot be fully removed from rs-fMRI data due to their spatial heterogeneity (61, 63). In fact, the four components identified in this work depict distinct spatial patterns which are also partially overlapped with each other. There has been a long history of debate on whether global signal regression should be applied to fMRI data (64). Instead of heuristically regressing out a single time course (the global signal) for the entire brain during the global signal regression, NASCAR is capable of modeling these spatially heterogeneous components as low-rank factors, hence perhaps providing an improved approach to decoupling neuronal brain networks from physiological components.

### Multi-dimensional resting-state network atlas and applications

By generating these highly reproducible, spatially overlapped, and temporally correlated brain networks, we have constructed a “functional network atlas”, representing the functional spatiotemporal organization of the human brain common across the normal (healthy) population. This atlas can be used to study group differences in brain networks and is particularly powerful for studies with a limited number of subjects/patients. Using this functional network atlas, one could obtain a reliable estimation of brain network properties by a least-square fitting of individual subject’s rs-fMRI data to the functional atlas, which could be difficult if the analysis is done directly on the small dataset.

The subject participation level found from rs-fMRI data using NASCAR can be viewed as a feature extraction procedure from a machine learning perspective. Most rs-fMRI methods designed for studying brain networks (e.g., seeded correlation, ICA, graph theory, etc.) do not directly provide features for classification/prediction. On the other hand, methods designed for classification/prediction (e.g., hand-crafted feature extraction with traditional classifier/regressor or deep-learning-based method) may lack interpretability of extracted features due to the high dimensionality of the feature space and black-box operations. In contrast, the tensor-based feature extraction strategy bridges these two issues as we have shown in the application to a study of ADHD study: the subject mode not only provides individual network information for classification and prediction of ADHD indices, but also allows us to investigate differences in brain activity between ADHD subjects and controls in a variety of brain networks concurrently, hence potentially facilitating the exploration of neurological differences associated with ADHD. Applications of the procedure used in the ADHD study to other neurological and psychological conditions could be promising future directions.

### Limitations

While BrainSync could help temporally align fMRI data between subjects, the tensor-based NASCAR pipeline requires accurate spatial inter-subject coregistration. The low-rank model will fail if there is substantial spatial misalignment between subjects, although importantly, the same concern arises with group-ICA and related decompositions. In this work, we rely on the boundary-based method (65) and the multi-model surface matching method (66) for fMRI-to-T1 and cross-subject registration used in the HCP minimal preprocessing pipeline (32). Since the finalization of the HCP protocol, many model-based (66, 67) and deep-learning-based approaches (68) have been proposed to improve inter-subject coregistration. The combination of BrainSync with these more advanced spatial coregistration approaches could be potentially beneficial for further improvement of the network identification results. A second potential limitation is that BrainSync will perfectly align time series only to the degree that the underlying temporal-correlation structure is common among subjects. The differences among subjects after alignment reflect underlying differences in this correlation. However, since we use an unbiased group-based approach in the BrainSync transform (29) to find the best match across all subjects, these differences can be viewed as informative indicators of the way in which each individual differs from the group. It is these differences that are encoded in the subject participation scores used in the ADHD study.

As with any other data-driven approach (e.g., ICA), manual inspection of the components is always required. In particular, we note that there is a sign ambiguity associated with the networks that affects the interpretability of the results. Specifically, flipping the sign of both the time series and spatial model produces an identical fit to the data ((−***a***) ∘ (−***b***) = ***a*** ∘ ***b*** in Eq. 1). In this study, we adjusted network polarity based on inspection of the four correlation matrices shown in Fig. 3. An advanced and perhaps automated method for network recognition or a model that could avoid this ambiguity is highly desired.

## Materials and Methods

### The Human Connectome Project dataset

We used the minimally preprocessed 3T resting-state fMRI (rs-fMRI) data from 1000 subjects in the publicly available Human Connectome Project (HCP) database (32, 33). Each subject has two sessions of scans with two different phase encoding directions (LR, RL). We used both sessions from LR direction only, as a substantial number of subjects underwent incorrect preprocessing steps in the RL runs (69). Also, using a single phase encoding direction could help minimize the potential inter-subject misalignment due to the different EPI distortions in different directions, although EPI distortion had been carefully corrected during the preprocessing (70). These data were acquired with *TR* = 720 ms, *TE* = 33.1 ms and a 2 mm^3^ isotropic resolution and co-registered onto a common atlas in MNI space. Each session ran 15 minutes, with 1200 frames in total. The data were resampled onto the cortical surface extracted from each subject’s T1-weighted MRI and co-registered to a common surface (32), where there are approximately 22*K* vertices across the two hemispheres. We note that no additional spatial smoothing was applied beyond the 2 mm full width half maximum isotropic Gaussian smoothing used in the minimal preprocessing pipeline (32) because linear smoothing can blur boundaries between functional regions (71, 72).

### The ADHD-200 datasets

We used all 259 subjects from Peking University in the ADHD-200 dataset (34, 42). We excluded subjects with incomplete evaluations or measures, such as ADHD index, based on their phenotypic information. The total number of subjects that satisfy the requirements was 221, where 95 of them were diagnosed as having ADHD and the remaining 126 were neurotypical (NT). The rs-fMRI data were recorded using a Siemens Trio 3T scanner with a *TR* = 2 s. We preprocessed the rs-fMRI data using our previously developed BrainSuite fMRI Pipeline (BFP) (73). After applying the BFP pipeline, the rs-fMRI data were co-registered and re-sampled onto the same tessellated surfaces as used in the HCP dataset. We identified eleven outliers based on the Cook’s distance metric: A linear regression model was fit to the ADHD index using the estimated subject participation level described below. Then the Cook’s distance was measured for each subject. Subjects who had a Cook’s distance higher than the standard threshold 4/*N* (*N* = 221, the total number of subjects) (74) were identified as outliers.

### Brain network identification pipeline

The blood-oxygen-level-dependent (BOLD) signals in rs-fMRI data from different subjects are not temporally locked, hence not directly comparable in time. However, the low-rank tensor model described below requires temporal synchrony across subjects. We used the BrainSync algorithm to achieve temporal alignment of the rs-fMRI data (28). BrainSync finds an optimal temporal orthogonal transformation between two subjects, such that the time series in homologous regions of the brain are highly correlated after synchronization. To avoid potential bias in the selection of the specific reference subject, we used the group version of the BrainSync algorithm (29) to build a virtual reference subject, which is closest, in the mean-square sense, to all subjects. We then temporally aligned all subjects’ data to the virtual reference to obtain a multi-subject synchronized dataset, Fig. S6 (a). Let ***X*** ∈ ℝ^*V*×*T*^ be the synchronized rs-fMRI data of an individual subject, where *V* ≈ 22*K* is the number of vertices (space) and *T* = 1200 is the number of time points (time). Then all subjects were concatenated along the third dimension (subject), forming a data tensor ***𝒳*** ∈ ℝ^*V*×*T*×*S*^, where *S* is the number of subjects, Fig. S6 (b). We model brain networks present in the group rs-fMRI data as a low-rank Canonical Polyadic (CP) model, Fig. S6 (c):

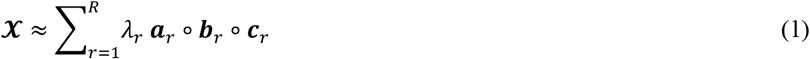

where each component *λ*_*i*_ ***a***_*i*_ ∘ ***b***_*i*_ ∘ ***c***_*i*_ represents a brain network; ***a***_*i*_ ∈ ℝ^*V*^, ***b***_*i*_ ∈ ℝ^*T*^, and ***c***_*i*_ ∈ ℝ are the spatial map, the temporal dynamics, and the subject participation level, respectively, in the *i*^*th*^ network, Fig. S6 (d); *λ*_*i*_ is the magnitude of that network, indicating the relative strength of the activity in that network to other networks across all subjects. Each of the components ***a***_*i*_, ***b***_*i*_, ***c***_*i*_ are normalized to unit length within the NASCAR algorithm. *R* is the desired maximum number of networks. *R* = 50 was used in this work as a reasonable upper bound based on the rank of rs-fMRI based on reports in the literature (5, 75). We computed the expansion in Eq. 1 using the Nadam-Accelerated SCAlable and Robust (NASCAR) canonical polyadic decomposition algorithm (30, 31). NASCAR employs an iterative method using low-rank solutions as part of the initializations when solving higher-rank problems. The robustness of the solutions and the scalability to a large dataset is substantially improved by using this warm start approach. Its performance relative to other network identification methods has been demonstrated in applications to both electroencephalography (EEG) data (76, 77) and task fMRI data (30, 31). We randomly divided the 1000 HCP subjects into two equally sized groups and applied this pipeline to each group and each session independently (2 groups ×2 sessions = 4 sessions in total).

### Post-NASCAR processing: spectrum estimation, network matching, and recognition

While correlations are unaffected by the orthogonal BrainSync transform, the virtual reference space does not itself provide time series that can be interpreted with respect to their dynamics or frequency content. Consequently, for visualization and spectral analysis, we applied the inverse-BrainSync transform of the temporal dynamics (***b***_*i*_) for each network to an arbitrary default subject (subject 100307 in the HCP dataset). The spectrum of the inversed time series was then estimated using the Welch method (36) with 24 50% overlapped Hamming windowed segments, each of length 100. As with other data-driven approaches (e.g., ICA) (78), we manually examined the NASCAR components based on their spatial maps, estimated spectra, and subject participation level. Seventeen subject-specific (one subject showed high participation level and others showed near-zero participation) or physiologically implausible “noise” components (noisy spatial maps or spectra) were identified at the higher ranks and removed from further consideration (examples were shown in our previous work (30)). To enable inter-session comparison, networks from all other sessions were matched to the first group’s first session using the Gale-Shapley algorithm (35). Because a one-to-one correspondence between NASCAR components in different sessions is not guaranteed, we further selected twenty-three highly reproducible components by thresholding the maximum of the inter-session correlations across all pairs of sessions at 0.9, as shown in Fig. S1 in the Supporting Information. To label the likely functionality of these networks, we computed the Pearson cross-correlation matrices using the spatial (***a***_***i***_), temporal (***b***_***i***_), subject mode (***c***_***i***_), and the estimated spectrum. By jointly visualizing the structures and patterns in these cross-correlation matrices, we manually clustered the networks into six functional categories and ordered them based on the cluster categories as shown in Fig. 3.

### Estimation of the subject participation level using the functional network atlas

The functional network atlas could be reformed into a set of bases by vectorizing the outer product of the spatial map and the temporal dynamics of each network. Specifically, let ***g***_*i*_ = *vec*(***a***_*i*_ ∘ ***b***_*i*_), where *vec*(·) is the vectorization operator. Then ***G*** = **[*g***_1_, ***g***_2_, …, ***g***_*R*_] ∈ ℝ^*V T* ×*R*^ forms the network atlas bases. Let ***Y*** ∈ ℝ^*V*×*T*^ be any rs-fMRI single-subject data independent from the atlas. Then the subject participation level for this subject can be estimated by a least-square fit: ***c*** = ***G***^†^***y*** ∈ ℝ^*R*×1^, where ***y*** = *vec*(***Y***) ∈ ℝ^*VT*×1^, ***G***^†^ is the pseudo inverse of ***G***.

## Data and code availability

The data used in this study are publicly available from the Human Connectome Project, Young Adult Study (https://www.humanconnectome.org/study/hcp-young-adult) and the International Neuroimaging Data-Sharing Initiative ADHD-200 dataset (https://fcon_1000.projects.nitrc.org/indi/adhd200).

For research purposes, we release the multi-dimensional functional atlas via Figshare (https://doi.org/10.6084/m9.figshare.c.5739128) and the open-access code is released at the GitHub repository (https://neuroimageusc.github.io/main)

## Acknowledgment

This work was supported by National Institutes of Health grants R01-EB009048, R01-EB026299, R01-NS089212, and K23-HD099309.

