## Supporting Information for "The intrinsic spatiotemporal organization of the human brain - A multi-dimensional functional network atlas"

### Supplementary Materials

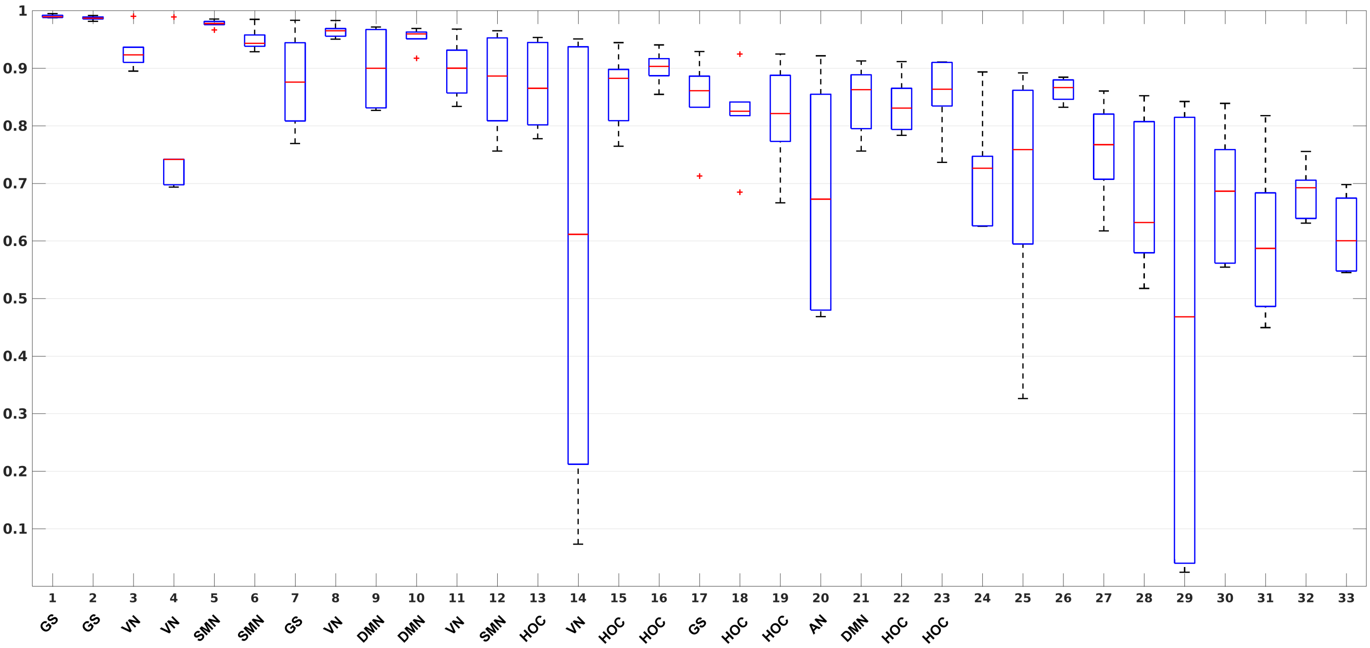


**Fig. S1**. Boxplots of the inter-session/inter-group Pearson correlations (across all 6 pairs of sessions)

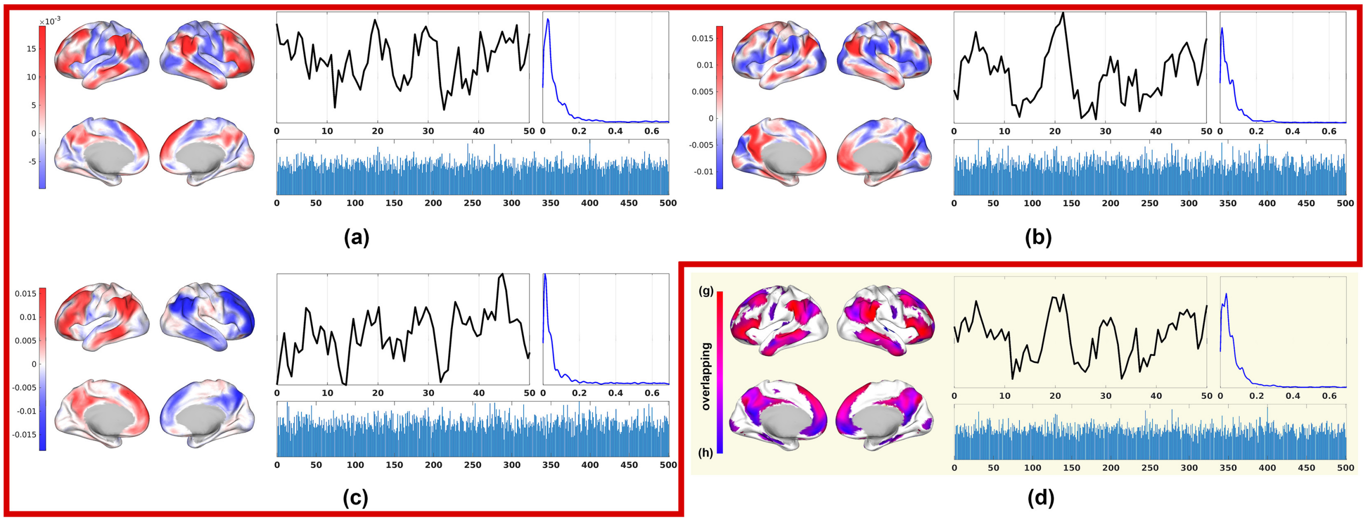


**Fig. S2.** Same set of brain networks as Fig. 1 but with inflated surface representations.

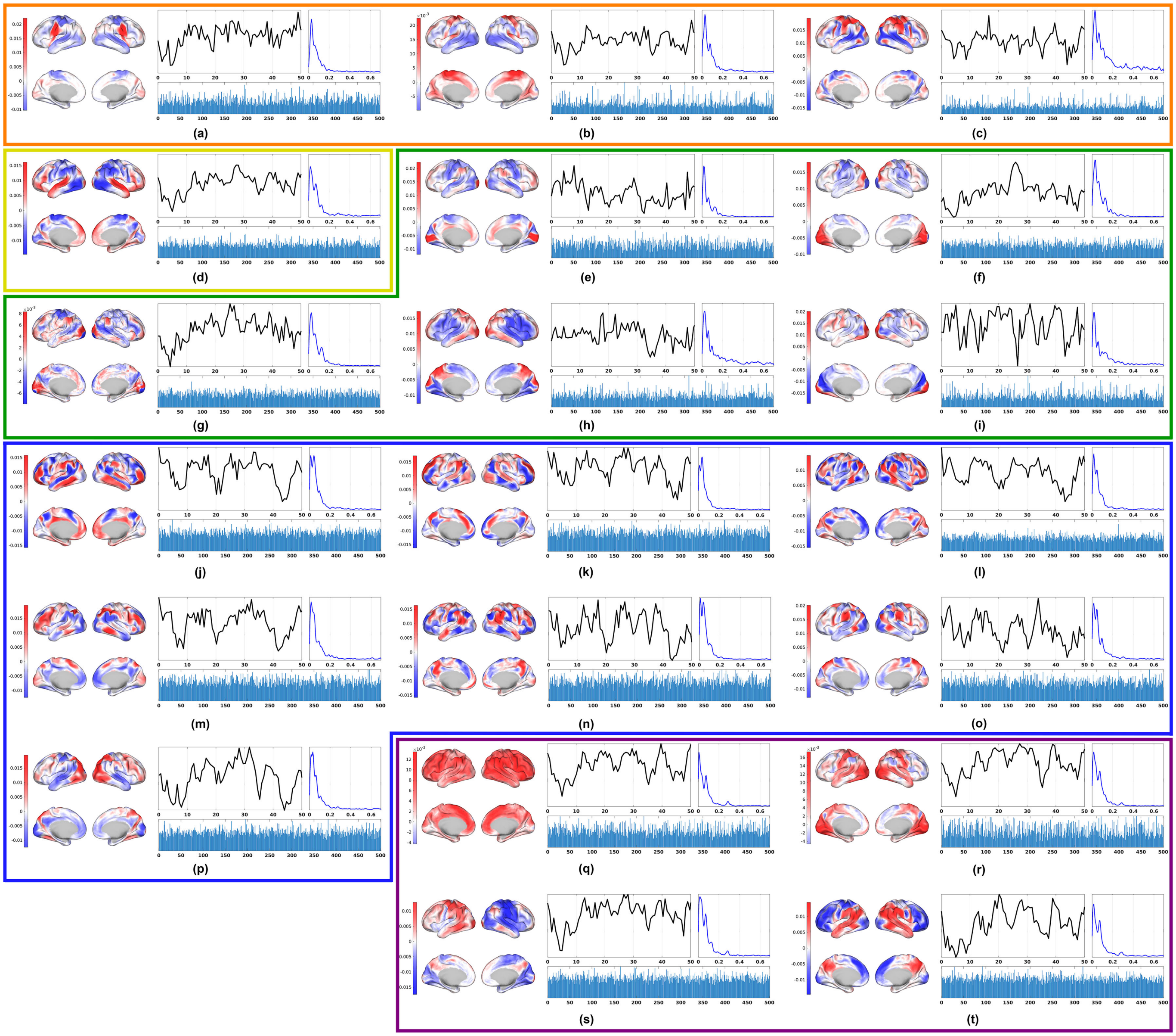


**Fig. S3.** Same set of brain networks as Fig. 2 but with inflated surface representations.

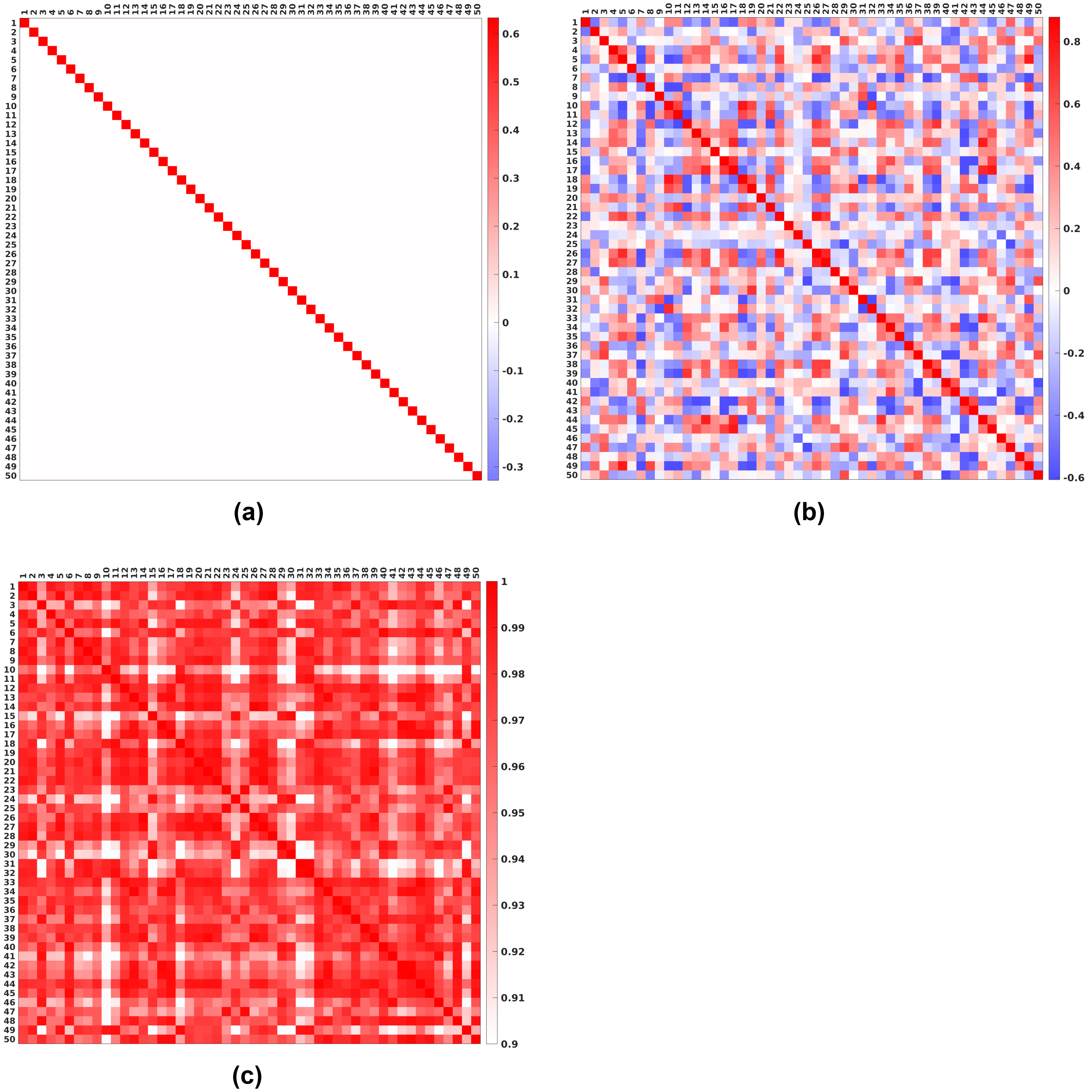


**Fig. S4.** Cross correlation matrices of the fifty networks identified using the group ICA method (Calhoun *et al.* 2001) and matched using the Gale-Shapley algorithm to the NASCAR results: (a) spatial correlation; (b) temporal correlation; (c) spectral correlation. Subplot (d) is omitted as the 2D-based ICA method does not provide subject participation information. The spectrum estimation and correlation computation processes were identical to that used in NASCAR and the color scales and color bars are also identical to those in Fig. 1.

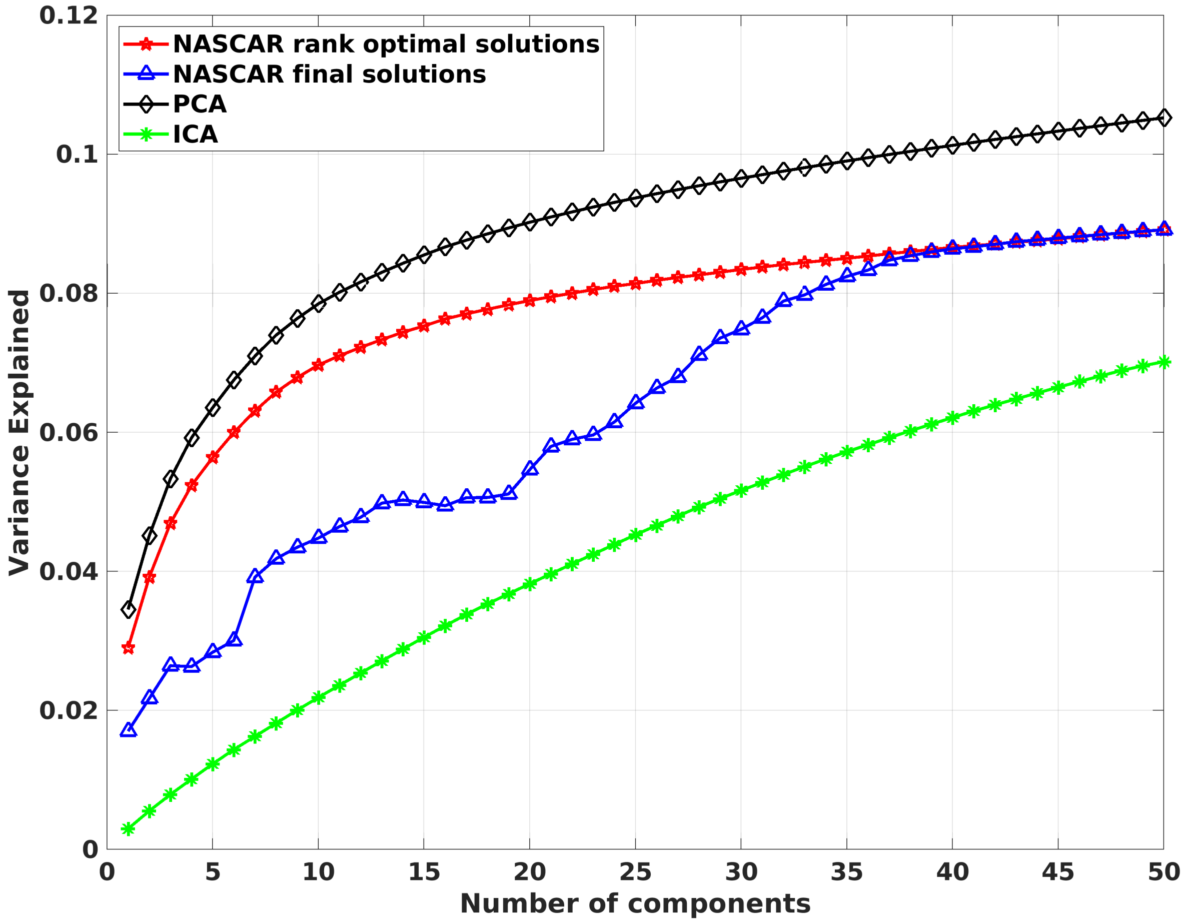


**Fig. S5.** Cumulative variance explained by NASCAR, PCA, and ICA. The red curve corresponds to the explained variance from the optimal NASCAR solutions at each rank, while the blue curve corresponds to the counterpart from the final NASCAR solutions.

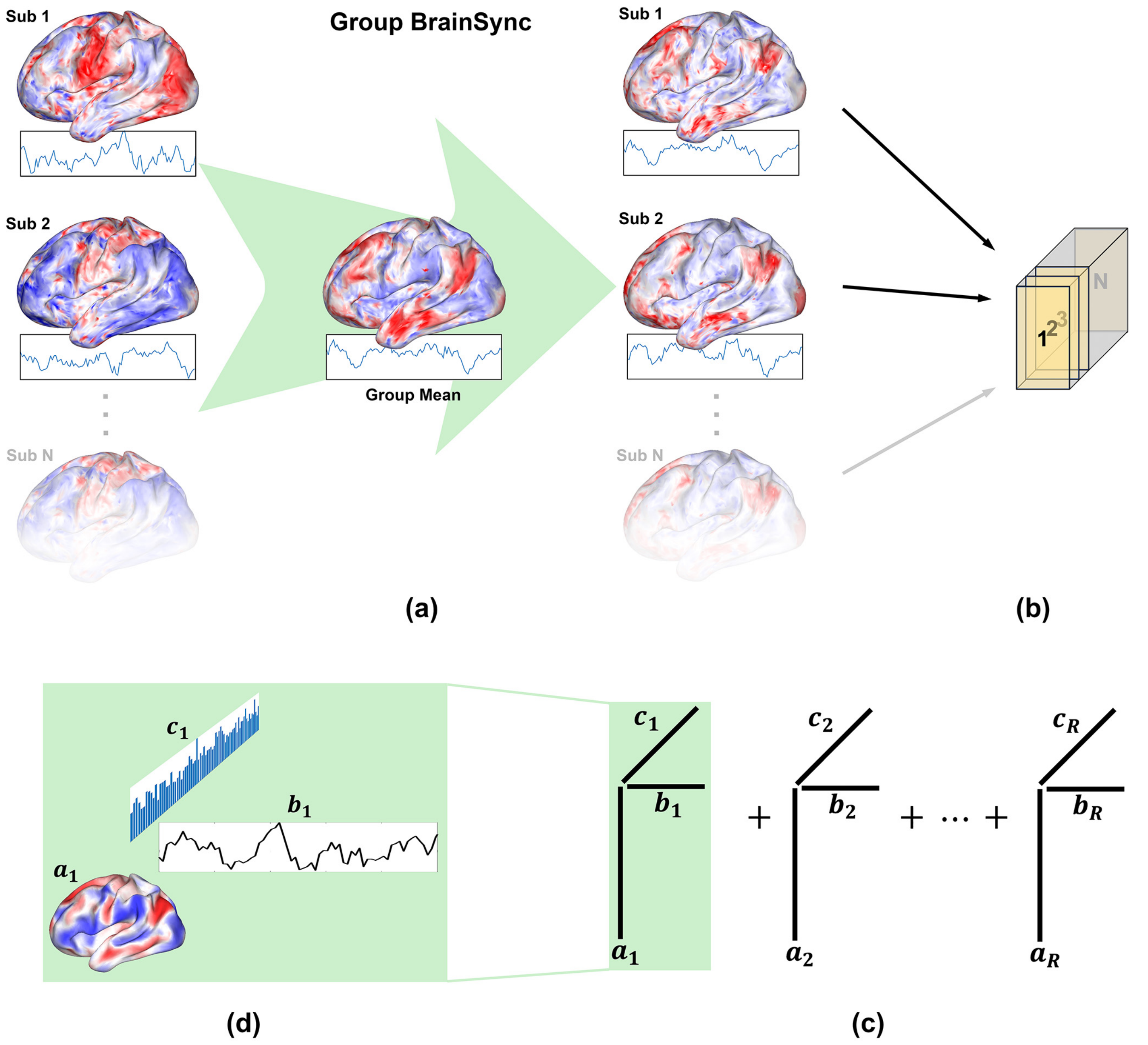


**Fig. S6**. Brain network identification from group rs-fMRI: (a) Resting-state data are temporally synchronized to a group mean using Group BrainSync; (b) a third-order tensor is formed with subject-index as the third dimension; (c) NASCAR tensor decomposition; (d) Visualization of the spatial map ($\boldsymbol{a}_i$), temporal dynamics ($\boldsymbol{b}_i$), and subject participation level ($\boldsymbol{c}_i$).

**Table S1.** Peak frequencies of each network/component

| Functional Category | Figure Index | 1st Peak Frequency (Hz) | 2nd Peak Frequency (Hz) | 3rd Peak Frequency (Hz) |
|---|---|---|---|---|
| Default mode networks | 1a | 0.027 | | |
| | 1b | 0.011 | | |
| | 1c | 0.011 | | |
| Somatomotor networks | 2a | 0.027 | | |
| | 2b | 0.027 | 0.076 | |
| | 2c | 0.027 | 0.065 | |
| Auditory network | 2d | 0.016 | 0.071 | 0.125 |
| Visual networks | 2e | 0.016 | 0.06 | 0.119 |
| | 2f | 0.027 | 0.06 | 0.125 |
| | 2g | 0.027 | 0.06 | 0.125 |
| | 2h | 0.022 | 0.06 | 0.098 |
| | 2i | 0.016 | 0.06 | 0.098 |
| Higher-order cognitive networks | 2j | 0.011 | 0.043 | |
| | 2k | 0.011 | 0.038 | |
| | 2l | 0.016 | 0.043 | |
| | 2m | 0.016 | | |
| | 2n | 0.011 | 0.038 | |
| | 2o | 0.016 | 0.043 | 0.092 |
| | 2p | 0.016 | 0.033 | |
| Global signals | 2q | 0.005 | 0.076 | 0.288 |
| | 2r | 0.016 | 0.076 | 0.288 |
| | 2s | 0.016 | 0.071 | 0.282 |
| | 2t | 0.016 | 0.071 | 0.288 |
